# Plant responses to wheat curl mites and wheat streak mosaic virus (WSMV): first evidence of virus tolerance in wheat

**DOI:** 10.1101/2024.09.15.612927

**Authors:** Lise Pingault, Tessa Albrecht, Kirk Broders, Jennifer Rushton, Joe Louis, Punya Nachappa, Vamsi Nalam

## Abstract

Wheat curl mites (WCM) are arthropod pests that pose significant threats to wheat crops by causing direct damage by feeding, and transmitting viruses such as Wheat Streak Mosaic Virus (WSMV), Triticum Mosaic Virus (TriMV), and High Plains Wheat Mosaic Virus (HPWMoV), leading to substantial losses in wheat, barley, oats, and rye. Over three years of field screening, we found that the cultivar Hatcher consistently produced higher yields under high WSMV disease pressure, outperforming Mace and TAM112, which carry the *Wsm1* gene and a QTL for curl mite resistance, respectively, indicating tolerance. To investigate the mechanisms underlying the tolerance phenotype in Hatcher, we compared its response to WCM and WSMV infection with a susceptible genotype, CO15D173R. Transcriptomic analysis revealed a nuanced interplay between plant defense and growth in Hatcher, with upregulation of genes related to jasmonic acid (JA), salicylic acid (SA), and abscisic acid (ABA) pathways, indicating a coordinated defense response. The activation of lignin biosynthesis genes points to a potential role of cell wall strengthening in deterring WCM and WSMV. Additionally, the regulation of genes involved in growth-related hormonal pathways such as gibberellic acid (GA), and brassinosteroids (BR) highlights Hatcher’s ability to maintain growth disease pressure. Our findings provide insight into the intricate network of phytohormones, growth-defense trade-offs, and cell wall modifications contributing to Hatcher’s tolerance to WCM and WSMV. This knowledge can inform the development of tolerant wheat varieties and enhance integrated pest management strategies, ultimately safeguarding wheat production.

## Introduction

Wheat curl mites (WCM; *Aceria tosichella* Keifer) are important arthropod pests of bread wheat (*Triticum aestivum* L.) worldwide. WCM cause direct damage by feeding, leading to reduction in yields (Harvey et al. 2000). More importantly, WCM-transmitted viruses, including the *Wheat streak mosaic virus* (WSMV; genus *Tritimovirus*, family *Potyviridae*) (Slykhuis 1955), *Triticum mosaic virus* (TriMV*; Poacevirus*, *Potyviridae*) (Seifers et al. 2009), and *High Plains wheat mosaic emaravirus* (HPWMoV; *Emaravirus*, *Fimoviridae*) (Seifer s et al. 1997), are among the most significant viruses impacting U.S. wheat production (Burrows et al. 2009; Navia et al. 2013; Murphy and Burrows 2021). Pests impacting wheat plants represent 10-50% of yield losses worldwide (Savary et al. 2019; Oerke 2006). The average yield losses from the WCM-WSMV complex range from 2-3% in the U.S. Great Plains, but up to 100% yield losses may occur in severely affected fields (Appel et al. 2015; Hollandbeck et al. 2020).

As there are no effective chemical solutions available to manage WCM (Murphy and Burrows 2021), the management of the complex virus-vector via plant breeding is critical to maintain wheat production. The management of WCM-WSMV disease complex has focused on an Integrated Pest Management (IPM) approach that combines controlling alternative hosts such as volunteer wheat, corn and wild grassy weeds, delayed planting to avoid any overlap between maturing summer crops and newly emerging winter wheat seedlings and use of mite and virus-resistant varieties [reviewed in (Nachappa et al. 2020; Singh et al. 2018; Skoracka et al. 2018; Tatineni and Hein 2018)]. So far, three (*Cmc2*, *Cmc3* and *Cmc4*) and four (*Wsm1*, *Wsm2*, *Wsm3* and *c2652*) loci conferring resistance to WCM and WSMV, respectively, have been characterized in the wheat genome (Malik et al. 2003; Zhao et al. 2021; Haber, Seifers, and Thomas 2006; Tatineni et al. 2016; Xie et al. 2022). Reports of virulent WCM populations and potential resistance-breaking WSMV isolates highlight the need for more diverse sources of resistance [Reviewed in (Nachappa, Haley, and Pearce 2021; Fellers et al. 2019; Albrecht et al. 2022; Redila, Prakash, and Nouri 2021))].

Various plant signaling pathways have been characterized in plants for tolerance to herbivory. Notably, phytohormones play a role in plant development, as well as response to biotic and abiotic stresses. Previously, it has been shown that jasmonic acid (JA) and abscisic acid (ABA) play key roles in providing tolerance to aphids in sorghum and soybean (Chapman et al. 2018; Grover et al. 2020). In addition to secondary metabolism compounds, plants utilize physical barriers to deter pest feeding. Lignin is one of the most abundant biopolymers in plants, which can enhance both the rigidity and thickness of the plant cell wall (Vanholme et al. 2008), making lignin an important defensive barrier against pests and diseases. Similarly, leaf morphology integrity is critical in response to WCM feeding. WCM feed on wheat epidermal tissues between the leaf veins and prevent the leaves to uncurl by damaging the bulliform cells (Skoracka, Rector, and Hein 2018). The whorl and curled leaves of the host plant provides a more humid micro-environment for WCM, which benefits WCM for their survival and reproduction, and shelter them from predatory and miticidal exposure. Thus, maintaining the integrity of the plant cell wall is critical to prevent the WCM from creating a beneficial micro-environment and for the plant to maintain the photosynthesis. Genes involved in wheat lignin biosynthesis pathway have been characterized (Nguyen et al. 2016) and multiple phytohormones have been shown playing a positive or negative role in the regulation of lignin biosynthesis, such as salicylic acid (SA), auxin (IAA) and cytokinin. An inverse relationship was observed for the SA level and lignin content (Gallego-Giraldo et al. 2011). On the other hand, a positive correlation has been reported between IAA and lignin content, and it was observed that the IAA signaling was more significant compared to the amount for lignin formation in wheat stem (Herrero, Esteban Carrasco, and Zapata 2014; Nguyen et al. 2016). However, cytokinin levels enhanced lignin biosynthesis in wheat (Herrero, Esteban Carrasco, and Zapata 2014; Nguyen et al. 2016).

In this study, we investigated wheat responses to WCM and WSMV in both a tolerant genotype (Hatcher, a commercial cultivar used in the Great Plains) and a susceptible genotype (CO15D173R) through field and transcriptomic approaches. By exploring the molecular mechanisms underlying plant-mite-virus interactions, this research aims to contribute to the development of wheat varieties with enhanced tolerance and support the implementation of effective management strategies to address this significant threat to wheat production.

## Methods

### Plant and mite colony maintenance

The two wheat genotypes, Hatcher (WCM-WSMV tolerant) and CO15D173R (WCM-WSMV susceptible) were grown in half-gallon pots with two or three plants per pot in Promix HP^©^ growing medium. The laboratory WCM colony of viruliferous (WSMV-infected) and non-viruliferous mites was initiated from field-collected symptomatic wheat leaf samples. The symptomatic leaves were examined under a dissecting microscope for the presence of WCMs. Leaf samples that contained WCMs were transferred to healthy plants of susceptible wheat genotype, Pronghorn at the Z1.4 (Zadoks, Chang, and Konzak 1974) or the four-leaf stage.

Colony maintenance involved transferring ≥ 10 mites either singly with a fine camel-hair brush or by placing a small section of WCM-infested wheat leaf onto healthy plants. Wheat plants were placed in insect cages (45 x 45 x 76 cm, BioQuip, Compton, CA), outfitted with a no-thrips insect screen. The colonies were maintained in a 16:8 hour light-dark cycle at a temperature of 23°C. To prevent cross-contamination, viruliferous and non-viruliferous colonies were maintained in separate rooms and regularly tested to verify the presence of WSMV and the designated mite genotypes.

### Field screening for WCM and WSMV resistance evaluation

The field plots were established at the USDA-ARS Central Great Plains research center near Akron, Colorado. This location was chosen as there are persistent populations of WCM as well as WSMV, HPWMoV, and TriMV that provide an ideal environment for screening resistance to curl-mite vectored viruses. The experiment was set up as a randomized complete block design with five winter wheat varieties randomized in each of the six experimental blocks. Each plot was 1.54 m wide and ranged in length from 4-4.6 m, with the exact length of each plot measured to calculate yield accurately. Wheat varieties used were Hatcher, Snowmass (carries *Wsm2* resistance gene), Mace (carries *Wsm1* resistance gene), TAM112 (carries a QTL for resistance to WCM), and Pronghorn as the susceptible check. To ensure elevated disease pressure, the experiment was planted adjacent to corn plots harboring WCM on Sept. 25, 2015, Sept. 15, 2016, and Sept. 15, 2017. Virus symptoms were assessed through visual symptoms and ELISA tests to detect the presence of WSMV, HPMoV, and TriMV in 2015 and 2016 and by quantitative PCR in 2017. Ten leaf samples were randomly collected from each variety in each plot to test for the presence of the virus. Disease incidence was calculated as the percentage of leaf samples in each plot that tested positive for WSMV by ELISA or qPCR. Plots were harvested in July of the following year, and the weight and moisture of kernels from each plot were recorded to calculate yield. We tested whether yield varied between varieties and years using a two-way ANOVA and completed multiple pairwise comparisons using Tukey’s HSD in R v. 4.2.2.

### Quantification of WSMV in wheat samples

Double antibody sandwich (DAS) – ELISA was used to determine the presence of WSMV in the field-collected samples in 2015 and 2016 as per the manufacturer’s protocol (Agdia Inc.®, Elkhart, IN). Briefly, leaf tissue (approximately 300 mg) was ground in a general extraction buffer at a 1:10 (wt/vol) ratio. Samples (100 μl/well) were loaded into WSMV-capture antibody-coated 96-well ELISA plates (Thermo Scientific, Inc.) and incubated at 37°C for 1 h. After rinsing, the WSMV conjugate antibody was added and incubated for 1 h. Plates were treated with PNP (p-nitrophenyl phosphate, 100 μl) and left in the dark for 1 h. Absorbance (at 405 nm) was measured after incubation, and samples with absorbance values ≥ two times the negative control were deemed positive.

Quantitative real-time PCR (qRT-PCR) was used to quantify WSMV in field-collected samples from 2017, as described previously (Albrecht et al. 2022). Briefly, total RNA was extracted from 40 mg leaf tissue using the Direct-zol RNA purification kit (Zymo Research, Irvine, CA) following the manufacturer’s instructions. The extracted RNA (∼50 ng) was then used in a qRT-PCR duplex assay (Price et al. 2010) with a TaqMan RNA-to-Ct 1-step kit (Applied Biosystems, ThermoFisher Scientific) on a QuantStudio3 Real-Time PCR system (Applied Biosystems) for WSMV detection and quantification. Reaction conditions were set for reverse transcription at 48°C for 30 min, followed by initial denaturation at 95°C for 10 min. Amplification then proceeded through 40 cycles of denaturation at 95°C for 15 s and annealing/extension at 60°C for 1 min. Samples with a Cq value below 34.9 for WSMV were considered positive. Absolute quantification was used to determine WSMV copy number. Standard curves were generated using ten-fold serial dilutions (ranging from 5 × 10^-13^ g to 5 × 10^-18^ g of DNA) of a 319 bp amplicon of the WSMV nuclear inclusion B (NIb) gene, which served as the reverse transcription quantitative PCR target. The primers used for WSMV amplification were: WSMV_ NIb_Fq (5’-CAAAGCTGTGGTTGATGAGTTCA-3’) and WSMV_NIb-Rq (5’-TTGATTCCGACAGTCCATG-3’) and WSMV_NIb_Rseq (5’-TCGAAACTTCTGCACAATCG-3’). The Cq values obtained from each dilution were plotted against the estimated copy number corresponding to each target DNA concentration. A linear regression was performed, ensuring an acceptable R^2^ value of > 0.9991.

### Sample collection, RNA extractions and sequencing

The experiment was a factorial design with two levels of the wheat variety factor (Hatcher and CO15D173R) and two levels of WCM andWSMV infestation (uninfested and infested). Each treatment combination was replicated three times. The two wheat varieties were germinated and vernalized for 10 weeks. Plants randomly assigned to the WCM and WSMV treatment were infested with viruliferous mites two weeks post-vernalization. For infestation, wheat leaf clips measuring approximately 2 cm and containing 15-20 viruliferous WCMs, were placed in the leaf whorl of the test plants. Wheat clippings are free of viruliferous WCMs and were laced inside the leaf whorls of plants assigned to the uninfested treatment. Both sets were placed in separate growth chambers to prevent contamination. Leaf tissue was collected from both sets of plants at the booting stage (Feeks 8-9) and immediately frozen in liquid nitrogen (LN2) and stored at −80 °C until further sample processing. From a subset of plants from this experiment, phenotypic measurements including the number heads, and tillers was also counted at weekly intervals until harvest.

Total RNA was isolated from 100 mg of frozen leaf tissue per sample using either Direct-zol® RNA Purification Kit or Quick-RNA^TM^ Miniprep Kit (Zymo Research, CA, USA), according to the manufacturer’s recommendations. RNA quality and quantity was assessed using the NanoDrop One spectrophotometer (Thermo Fisher Scien-tific, Waltham, MA) and Agilent 2100 Bioanalyzer (Santa Clara, CA). The plant RNA samples were sent to Novogene Corporation Inc. (Sacramento, CA) for cDNA library construction and RNA sequencing. The poly(A) mRNA-enriched libraries were constructed using NEBNext® Ultra™ II RNA Library Prep Kit (New England BioLabs, Ipswich, MA). Paired-end sequencing (2×150 bp) was conducted with the Illumina HiSeq 2000 platform. The raw sequencing reads have been deposited in the NCBI SRA database under the BioProject accession, PRJNA998910.

### RNA-seq analysis

The quality of the mRNA-seq (RNA-sequencing) raw reads was investigated with FastQC (Andrews 2010) and reads were trimmed with Trimmomatic using the following parameters: LEADING:20 TRAILING:20 SLIDINGWINDOW:4:20 MINLEN:75 (Bolger, Lohse, and Usadel 2014). Trimmed reads were mapped on the wheat reference genome v2.1 (Zhu et al. 2021). Tophat v2.1.1 (Kim et al. 2013) was used for the mapping using 2 mismatch, fr-firststrand library and unique mapped reads parameters (-N 2 –M). The transcripts reconstruction was performed with Cufflinks v2.2.1 with the following parameters: quantification against the reference annotation only (-G), multi-read-correct (-u) and frag-bias-correct (-b). The differential expressed gene analysis was performed with Cuffdiff 2.2.1. Differential expressed genes (DEGs) were identified with the following parameters: *q*-value ≤ 0.05 and fold-change (FC) |log_2_(Infested/Contol)| ≥ log_2_(2). Co-expression modules were identified using the weighted gene co-expression network (WGCNA) (Langfelder and Horvath 2008). Genes ID for the TFs were obtained from the PlantTFDB (http://planttfdb.gao-lab.org/index.php?sp=Sbi) using the reference genome v3.1. The GOBU package was used for enrichment calculations (Lin et al. 2006). The full set of wheat gene annotation was used as the reference comparison set against down or upregulated DEGs. The *P*-values were calculated using Fisher’s exact test and corrected for multiple testing with FDR method using the R module called ‘*p*-adjust’.

## Results

### Hatcher is tolerant to WCM and WSMV infection

WCM and WSMV was prevalent across the experimental site in each plot of each variety in all three years. Mean disease incidence of WSMV detected by ELISA revealed moderate to high disease incidence (25-93%) in 2015 and 2016 (Figure 1A). Pronghorn and Hatcher exhibited consistently higher WSMV incidences, while Mace and Snowmass showed significantly lower disease incidence in both years. In 2017, WSMV quantification using qRT-PCR showed less variation between varieties, with only Mace having a significantly lower WSMV load compared to the others (Figure 1B). The high WSMV copy number observed in 2017 and the high disease incidence in 2015 and 2016 suggest consistent WSMV pressure across all varieties and replicates throughout the study. Interestingly, despite the high disease pressure observed during the duration of the study, a significant interaction effect of variety, year, and variety-by-year interaction on yield was observed (Table 1). Hatcher, Snowmass, and Mace consistently produced higher yields in 2015 and 2016, while no significant differences were observed among varieties in 2017. When averaged across the three years, Hatcher had the highest overall yield, performing similarly to Snowmass (carries *Wsm2*). Hatcher significantly outyielded Mace (carries *Wsm1*) and TAM 112 (possessing a QTL for curl mite resistance) (Supplemental Figure 1).

**Figure 1.**
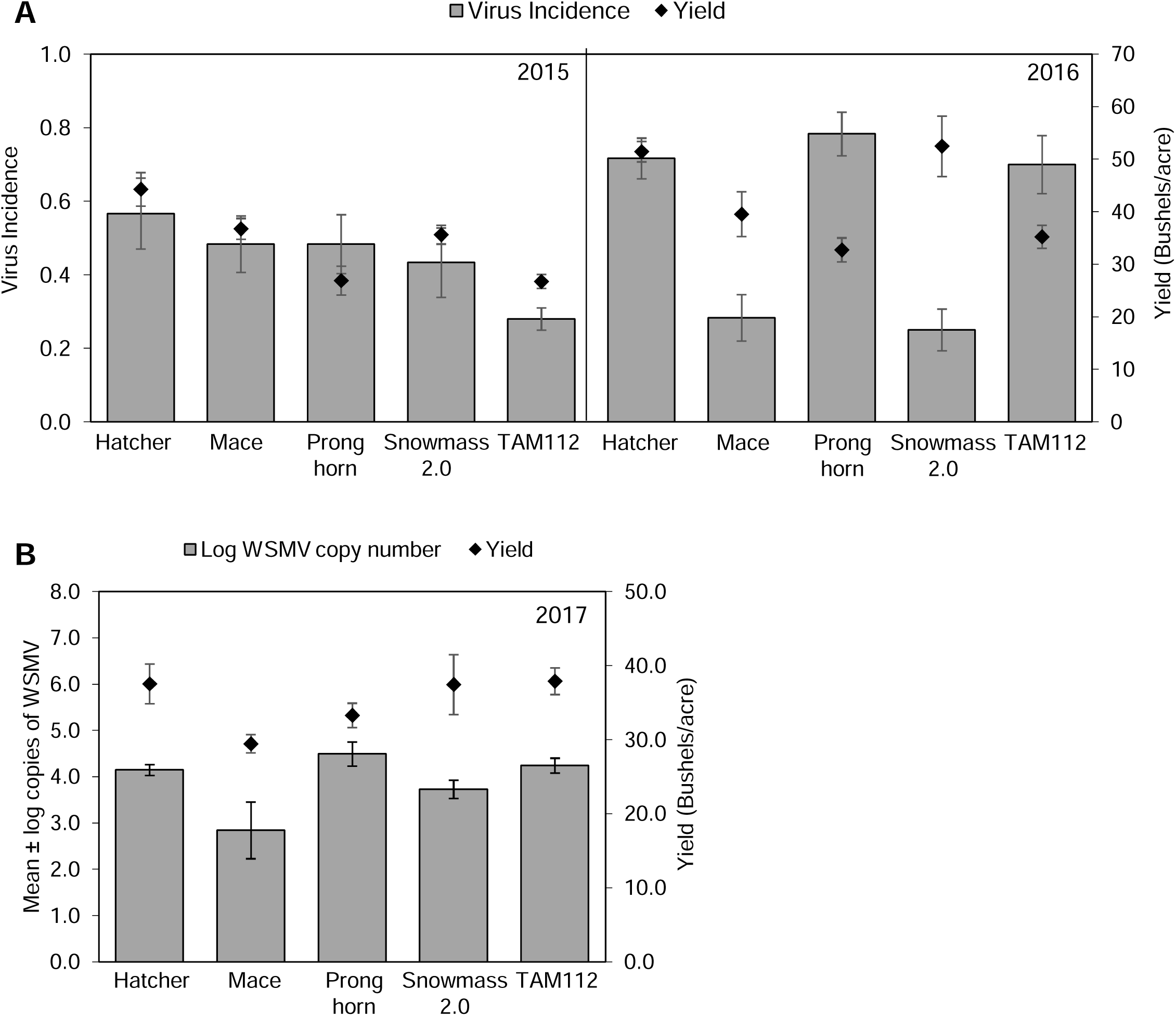
Hatcher is tolerant to WCM and WSMV infection. Response of wheat varieties to natural infection of wheat streak mosaic virus (WSMV) and wheat curl mites (WCM) in irrigated field trials conducted in (A) 2015, (B) 2016 and (C) 2017 (Akron, CO). In A and B, the bars indicate virus incidence and in C bars indicate mean of five biological replicates ± SEM log copies of WSMV RNA per variety. In A, B and C diamonds indicate average yield in bushels/acre.

**Table 1.** Analysis of Variance (ANOVA) for Wheat Yield across Varieties and Years.

|  | **Df** | **Sum Sq** | **Mean Sq** | **F value** | **Pr(>F)** |
| --- | --- | --- | --- | --- | --- |
| Variety | 4 | 2165 | 541.2 | 9.121 | 5.28e-06 |
| Year | 2 | 1199 | 599.3 | 10.1 | 0.000138 |
| Variety: Year | 8 | 1404 | 175.5 | 2.958 | 0.0065 |
| Residuals | 71 | 4213 | 59.3 |  |  |

In response to WCM and WSMV infection, the average number of heads and tillers were compared between Hatcher and CO15D173R over a10-to 70-day timecourse in a greenhouse experiment (Supplemental Figure 2). In uninfested plants, the number of head increased from 55 to 69 days, while infested plants showed significant difference in the number of heads across time points for both Hatcher and CO15D173R (Supplemental Figure 2A). Comparing infested and unifested plants within each genotype, the number of tiller was significantly lower in infested CO15D173R plants (*p*-value=0.04), whereas Hatcher showed no significant reduction in tillers (*p*-value=0.16) (Supplemental Figure 2B). Taken together, our data suggests that Hatcher’s ability to tolerate WCM and WSMV infection is part due to its ability to maintain growth despite disease pressure.

### Transcriptomic responses of wheat to WCM and WSMV

RNA-sequencing was performed on two wheat genotypes: Hatcher (tolerant to WCM and WSMV in our study, Figure 1) and CO15D173R (susceptible). Wheat plants were infested with viruliferous mites, and tissue samples were collected at the booting stage. Uninfested plants served as the control. Sequencing generated an average of 305 million paired-end reads per library, with 92% mapping to the hexaploid wheat reference genome (Supplemental Table 1). Analysis revealed the expression of 107,197 protein-coding genes across all conditions (Supplemental Table 2). Principal component analysis (PCA) was used to visualize the overall gene expression patterns across the four treatments (Figure 1, Figure 2A). The PCA plot indicates that genotype was the major factor influencing gene expression, with a clear separation between Hatcher and CO15D173R. Interestingly, even within each genotype, control (uninfested) and infested treatments clustered separately, suggesting an additional effect of WCM and WSMV on gene expression (Figure 1, Figure 2A).

**Figure 2.**
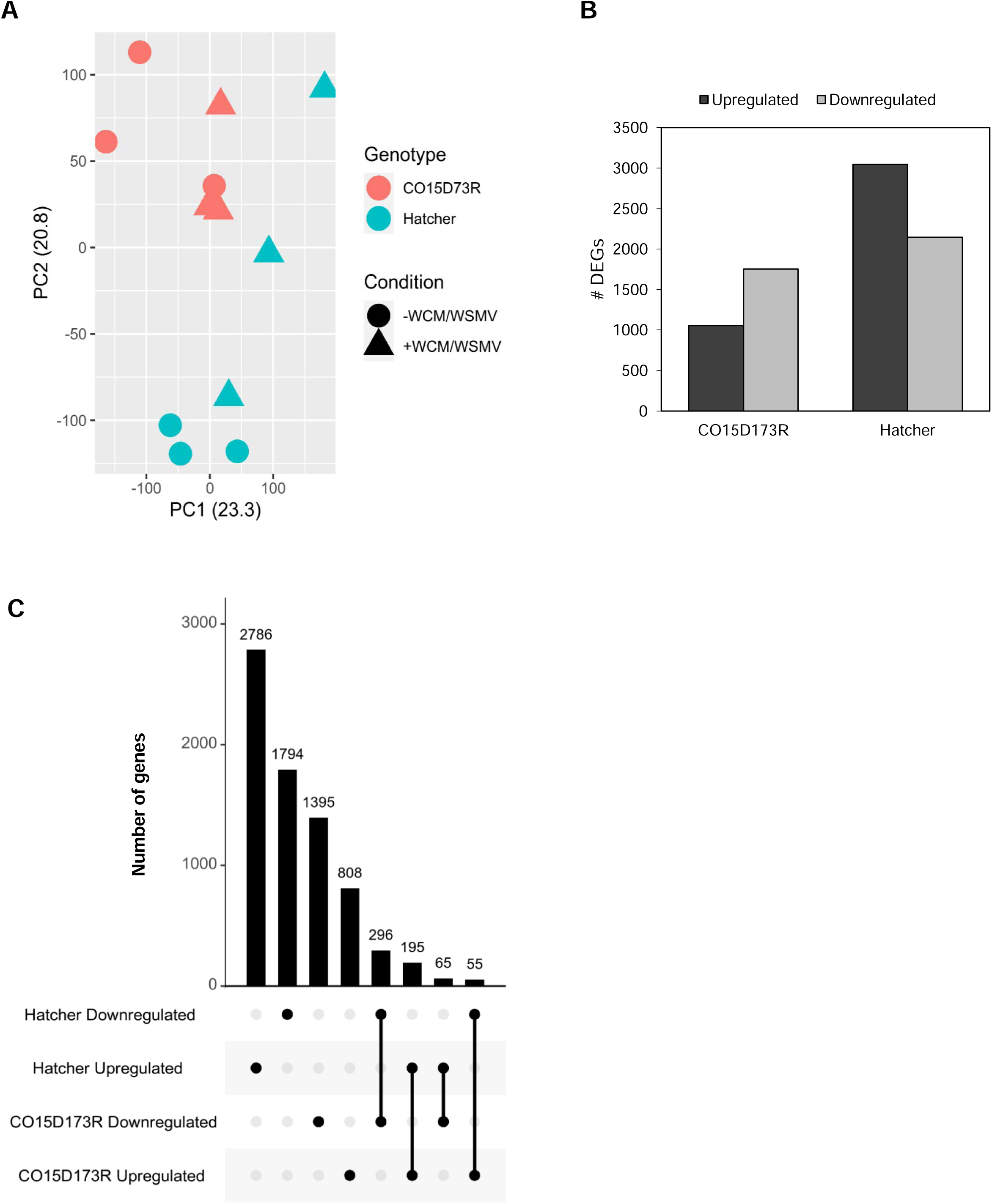
Wheat genomic response to WCM and WSMV. (A) PCA analysis of the 107,197 protein coding genes expressed in at least one condition. Wheat genotypes are represented with different colors (red= CO15173R and blue= Hatcher) and treatment condition with different shapes (circles = uninfested, triangles = Infested with WCM-WSMV). (B) The total number of genes up-or downregulated in each wheat genotype. (C) Visualization of the overlap of up or downregulated genes in the two wheat genotypes.

Differential gene expression analysis identified a total of 7,394 non-redundant DEGs responding to WCM and WSMV infestation, with a higher number observed in Hatcher (5,191 DEGs) compared to CO15D173R (2,814 DEGs) (Figure 2B). Notably, the susceptible CO15D173R exhibited a higher number of downregulated genes, while the tolerant Hatcher displayed a predominance of upregulated genes (χ^2^ = 324.58, *p*-value = 9.999e-05) (Figure 2B). This suggests that Hatcher’s tolerance may involve the activation of specific genes and pathways in response to infestation by viruliferous WCMs.

The UpSet plot approach identified DEGs common between multiple conditions (Figure 2C). Interestingly, no genes were identified that displayed differential expression in more than two conditions. The largest number of DEGs (2,786 genes) were exclusively upregulated in Hatcher, while the smallest subset was observed in CO15D173R (808 genes) (Figure 2B). Among the 611 DEGs between the conditions, 65 were upregulated in Hatcher but downregulated in CO15D173R, potentially representing key genes involved in tolerance to WCM and WSMV. Conversely, 55 genes were downregulated in Hatcher and upregulated in CO15D173R (Figure 2). While we did not find WCM resistance genes (*Cmc*) genes expressed in both of the transcriptomes, two candidate genes within the *Wsm2* locus of the reference genome (Xie et al. 2022) were upregulated in Hatcher: *Traes5A03G0661300* and *TraesCS5B03G0699800*, both encoding Chaperone protein DnaK. Additionally, a gene homologous to the Arabidopsis *RPM1* (*TraesCS3B03G0082600*) was downregulated in Hatcher and exhibited no differential expression in CO15D173R (Supplemental Table 2).

### GO analysis indicates functional differences in response to WCM and WSMV infestation

Gene Ontology (GO) enrichment analysis was performed to identify biological processes and molecular functions associated with DEGs in both wheat genotypes following WCM AND WSMV infestation. Upregulated genes in the GO-molecular function category in the tolerant Hatcher (2,786 genes) were enriched for functions potentially involved in defense responses (Figure 3A). These included phenylalanine ammonia-lyase (PAL) activity, monooxygenase activity, UDP-glycosyltransferase activity, and active membrane transporter activity for metabolite movement. Additionally, upregulated genes included those with calcium ion binding activity. Downregulated genes in Hatcher (1,794 genes) associated with GO-molecular function were enriched for functions related to photosynthesis, including chlorophyll-binding, oxidoreductase activity, and magnesium protoporphyrin IX methyltransferase activity (Supplemental Figure 3).

**Figure 3:**
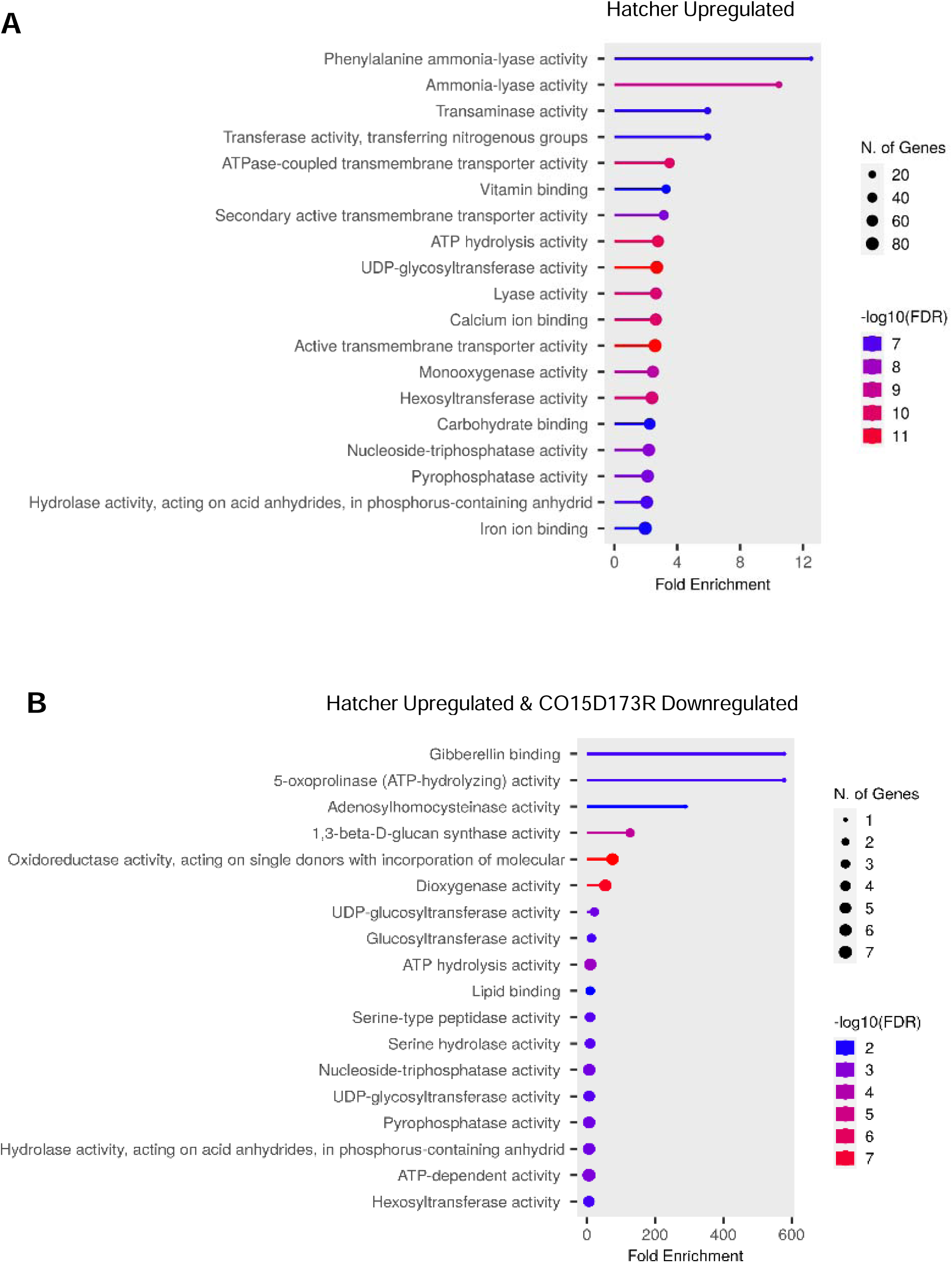
Comparison of functional enrichment analysis of the DEGs in wheat genotypes in response to WCM and WSMV infestation. (A) Hatcher upregulated genes after WCM **and** WSMV infestation (2,786 genes). (B) DEGs up or downregulated in Hatcher and CO15D173R (65 genes).

Upregulated genes in the susceptible genotype CO15D173R (808 genes) were enriched for functions not directly related to defense. These included structural molecule activity, protein transporting ATPase activity, and protochlorophyllide reductase activity, potentially associated with basic cellular processes. Downregulated genes in CO15D173R (1,395 genes) in the susceptible genotype were enriched for functions crucial for photosynthesis, including actin binding (crucial for chloroplast structure), chlorophyll-binding, and electron transport within the cyclic electron transport pathway of photosynthesis (Figure 3B). Additionally, the downregulation of genes related to 5’-3’ RNA polymerase activity and hydrolase activity might indicate a broader disruption of cellular processes. A small set of genes (195) displayed differential expression in both genotypes and were enriched for functions potentially involved in general stress responses. These included chitin binding, important for recognizing fungal pathogens, glutathione transferase activity involved in detoxification, glutamate dehydrogenase activity related to nitrogen metabolism, and general hydrolase activity. This analysis suggests that Hatcher, the tolerant genotype, may activate specific defense-related pathways in response to WCM and WSMV, while CO15D173R, the susceptible genotype, experiences disruptions in core cellular processes, including photosynthesis. The shared responses in both genotypes highlight common stress responses triggered by WCM and WSMV infestation.

### Metabolic pathway analysis reveals differences in response to WCM and WSMV infestation

MapMan (Thimm et al. 2004) was used to characterize the key pathways affected by WCM and WSMV infestation (Figure 4). Analysis of pathways involved in general metabolism revealed contrasting responses in key metabolic pathways between Hatcher and CO15D173R (Figure 4A, B). Upregulated genes in Hatcher were associated with processes like cell wall biosynthesis, indicating potential reinforcement against WCM and WSMV infestation. Conversely, downregulated genes in the susceptible genotype were enriched for pathways like photorespiration, suggesting a potential reduction in photosynthetic efficiency. For genes specifically involved in pathways related to biotic response, more downregulation was observed in the CO15D173R as compared to Hatcher (Figure 4A, B). Hormone signaling pathways, such as JA and ABA, were upregulated in the susceptible genotype. Additionally, genes encoding Ethylene Responsive Factor (ERF), transcription factirs involved in stress responses, were upregulated in the susceptible genotype and downregulated in the tolerant genotype (Figure 4A, B). In contrast, Hatcher displayed upregulation of genes involved in cell wall reinforcement, β-glucanase activity (associated with defense against fungal pathogens), heat shock proteins (involved in stress tolerance), and redox regulation (important for maintaining cellular balance). These genes were downregulated in CO15D173R (Figure 4A).

**Figure 4:**
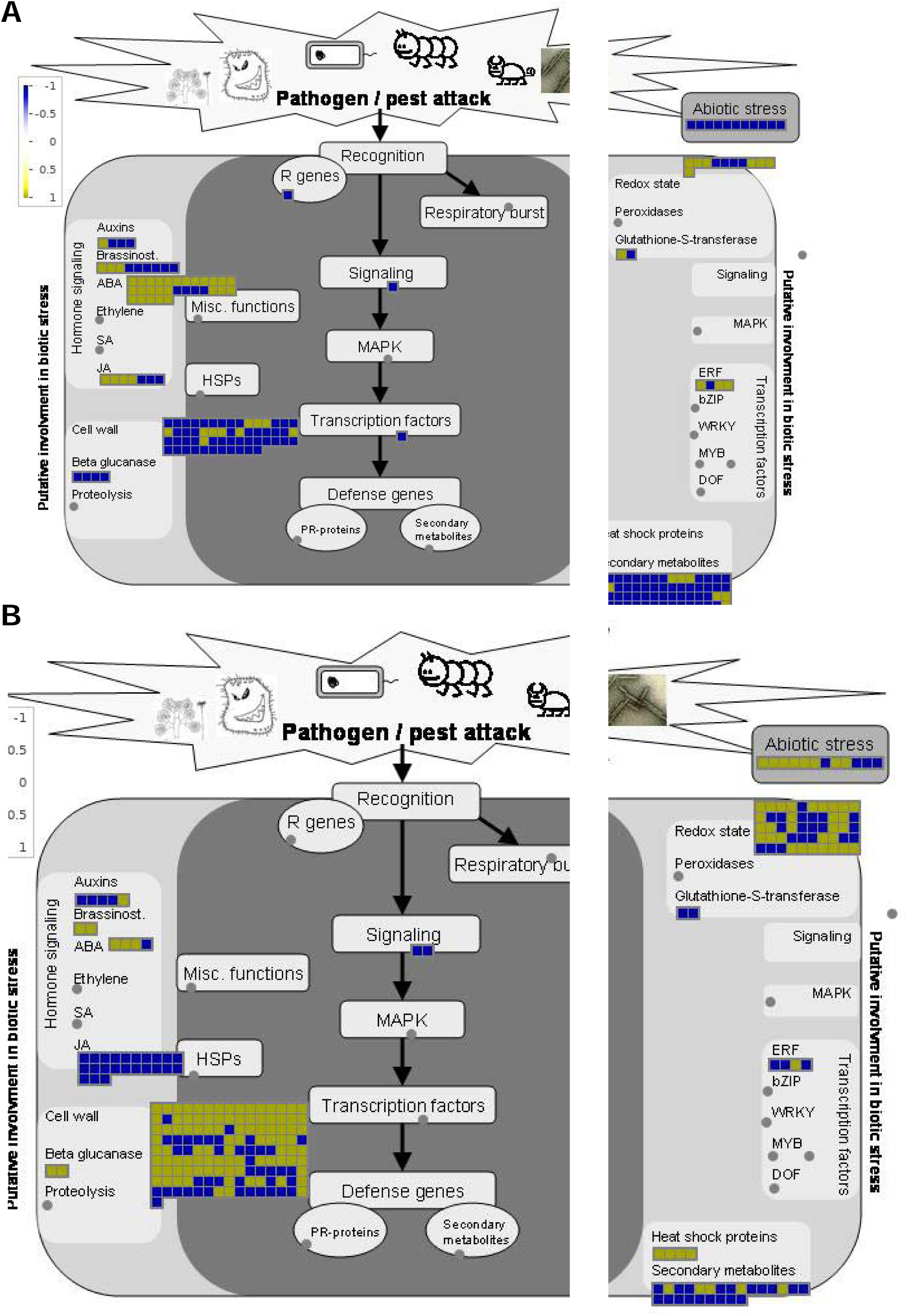
Overview of the transcriptomic response after WCM and WSMV infestation. Using MapMan representation of the gene expression in the susceptible genotype (A) and tolerant genotype (B). Each box represents the log_10_(FC). Yellow indicates upregulated gene expression and blue downregulated gene expression in response to WCM and WSMV.

Lignin is a complex phenolic polymer that forms an important structural component of the plant’s secondary cell wall. Among the 221 genes identified in the wheat lignin biosynthesis pathway (Figure 5 A; Nguyen et al. 2016), 61 were differentially expressed in response to WSMV+WCM infestation (Figure 5). Notably, among these genes 50 were found upregulated in Hatcher, encompassing all the major steps of lignin biosynthesis. Specifically, genes involved in converting the monolignol, *p-*coumaroyl-CoA to oligolignols and lignin via the shikimic acid pathway were all upregulated in Hatcher during WCM and WSMV infestation (Figure 5B).

**Figure 5:**
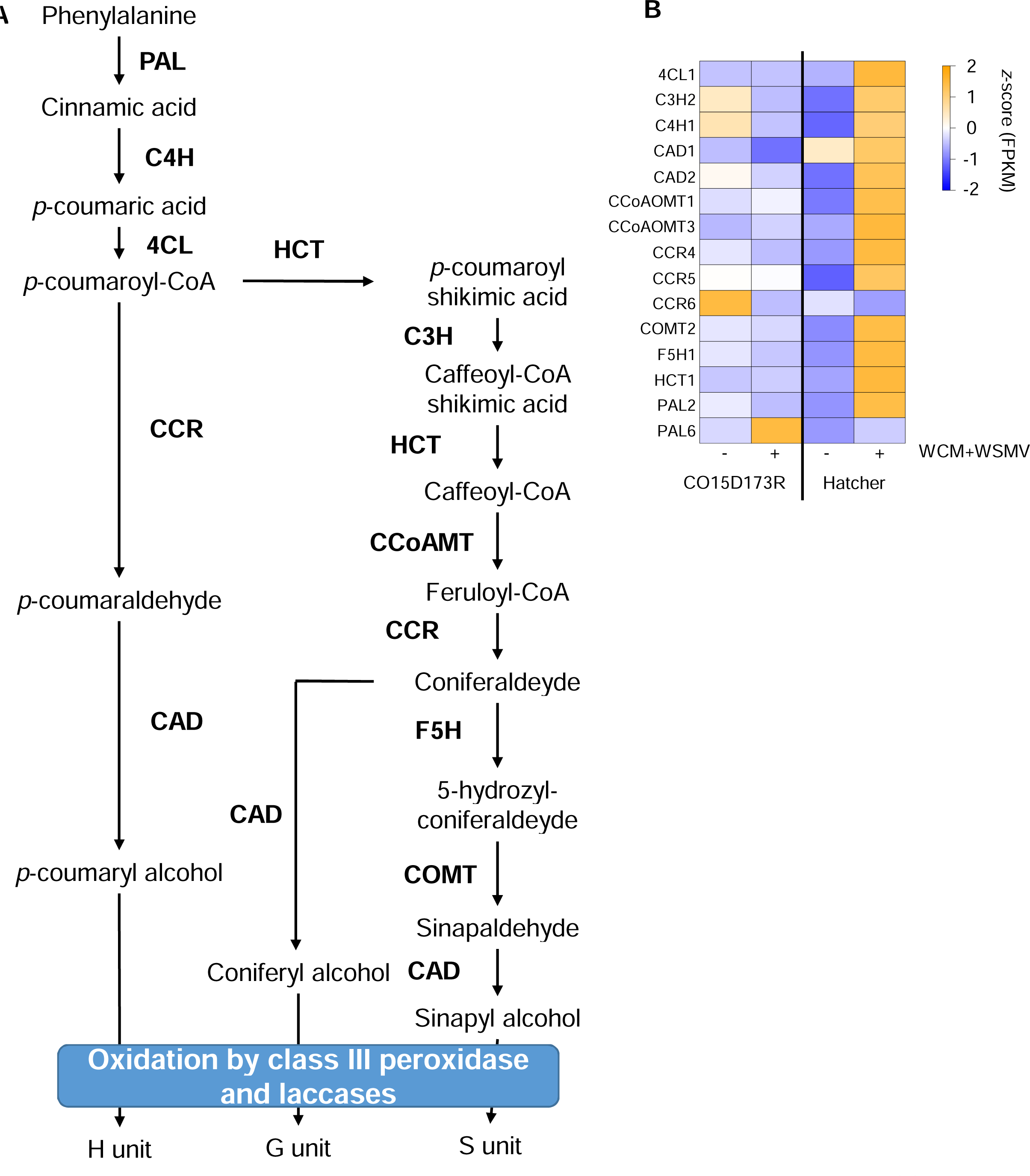
Overview of the transcription profile of genes involved lignin biosynthesis pathway. (A) Wheat lignin biosynthesis pathway adapted from Nguyen et al., 2016. (B) Heatmap of the expression of WCM and WSMV infestation induced change of genes encoding for lignin biosynthesis pathway. Expression values have been averaged for each gene family and normalized (*z*-score). The gene IDs and related FPKM values are indicated in the Supplemental Table 2.

### KEGG Pathway Analysis Reveals Disruptions in CO15D173R

KEGG pathway analysis provided further insights into the effects of WCM and WSMV infestation on various metabolic processes (Figure 6). Notably, genes downregulated in CO15D173R were enriched for functions related to core metabolic pathways, including carbohydrate, energy, lipid, amino acid metabolism, hormone biosynthesis, and signaling. This suggests a potential disruption of basic cellular processes in CO15D173R in response to WCM and WSMV infestation.

**Figure 6:**
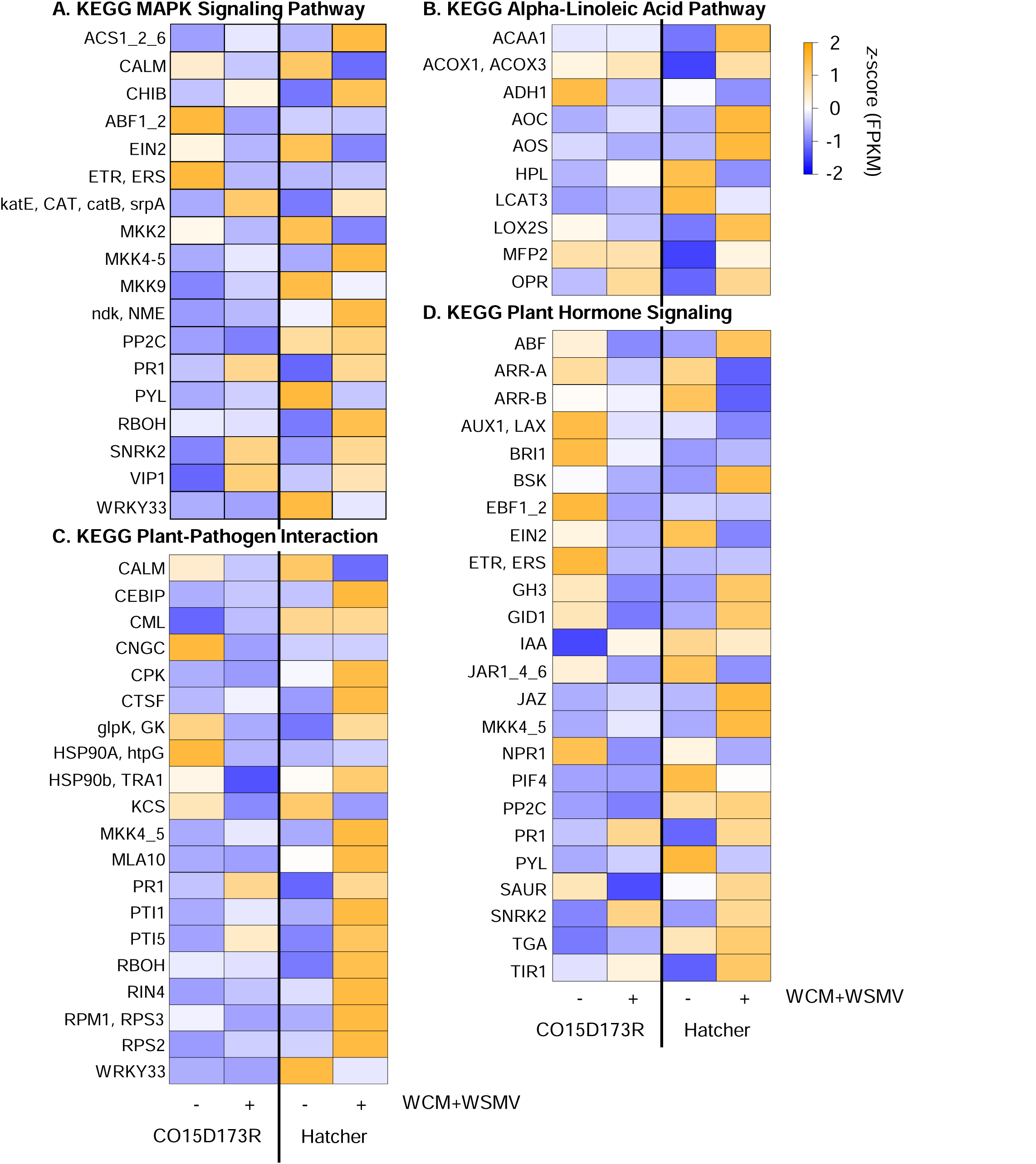
Heatmap of the expression of WCM and WSMV infestation induced change of genes encoding. (A) MAPK signaling pathway, (B) alpha-linoleic acid, (C) hormone signaling, and (D) plant-pathogen interaction pathways. Each column corresponds to a wheat genotype. Each cell contains the corresponding average *z*-score (FPKM) for each gene. Highly expressed genes are represented in yellow and low expressed genes in blue. The gene IDs and related FPKM values are indicated in the Supplemental Table 2.

Contrasting responses in hormone signaling pathways between Hatcher and CO15D173R were observed. Genes related to auxin response, *AUX1*, *GH3*, and *SAUR*, known to influence cell enlargement and plant growth, were upregulated in Hatcher but downregulated in CO15D173R. Similarly, genes involved in cytokinin response, associated with cell division and shoot initiation, were downregulated in Hatcher and CO15D173R. Interestingly, genes encoding gibberellin TF (*PIF3* and *PIF4*) were upregulated in Hatcher and CO15D173R. Ethylene signaling also differed between the two genotypes. Genes that are negatively regulated by ethylene, ETR, and EBF1/2, which in turn negatively regulates *EIN3*, were downregulated in CO15D173R, potentially indicating a dampened response. Conversely, genes induced by SA, a key defense hormone, were downregulated in the susceptible genotype but were highly upregulated in the tolerant genotype, suggesting a role for SA-mediated responses in Hatcher.

The expression of *NPR1*, which mediates SA signaling, was downregulated in the susceptible plant. Genes related to the α-linoleic acid pathway, which form precursors of JA, were upregulated in Hatcher, suggesting increased JA synthesis could possibly contribute to Hatcher’s tolerant phenotype (Figure 6B). Conversely, the susceptible genotype exhibited minimal upregulation and some downregulation of genes within this pathway (Figure 6B). Additionally, genes encoding functional pathogenesis-related proteins were downregulated in the susceptible genotype but were upregulated in Hatcher. The PYL genes, a family of PYR/PYL type ABA receptors, were downregulated in Hatcher but upregulated in CO15D173R (Figure 6C).

A suite of other pathways associated with plant defense responses also showed differential expression. For instance, 55 genes associated with MAPK signaling showed differential expression in both genotypes (Figure 6D). Notably, genes encoding a basic endochitinase B, enzymes that cleave chitin randomly, were upregulated in both genotypes, with a higher fold-change in Hatcher, suggesting the activation of chitin-based defense mechanisms. Seven other genes, *CNGCs*, *CDPK*, *CaMCML*, *NHO1*, *RPM1*, *HSP90*, and *KCS1/10,* associated with defense-related responses, specifically the suppression of defense gene expression, and hypersensitivity response (HR), were downregulated in the susceptible genotype (Figure 6D).

### Hierarchical Clustering Reveals Genotype-Specific Responses

The 7,394 DEGs were grouped into nine clusters based on their expression profiles across all four conditions (susceptible control, susceptible infested, tolerant control, and tolerant infested) (Figure 7). Two clusters, 4 and 5, contain genes with high expression in control conditions for Hatcher and CO15D173R, respectively (Figure 7). Two clusters, 1 and 6, were mainly associated with CO15D173R. Cluster 1 grouped genes were highly expressed in infested susceptible plants, potentially reflecting stress responses specific to this genotype. Cluster 6, on the other hand, included genes with high expression in both control and WCM and WSMV-infested conditions in CO15D173R, suggesting a potential lack of response regulation. Cluster 9 includes genes expressed in low levels in control condition for Hatcher and CO15D173R infested but with high expression in the two other conditions. These genes represent functions related to gibberellin binding, lipid binding or polysaccharide binding. These functions were activated after WCM and WSMV infestation in the tolerant plants, while their expression was reduced in the susceptible plants during infestation.

**Figure 7:**
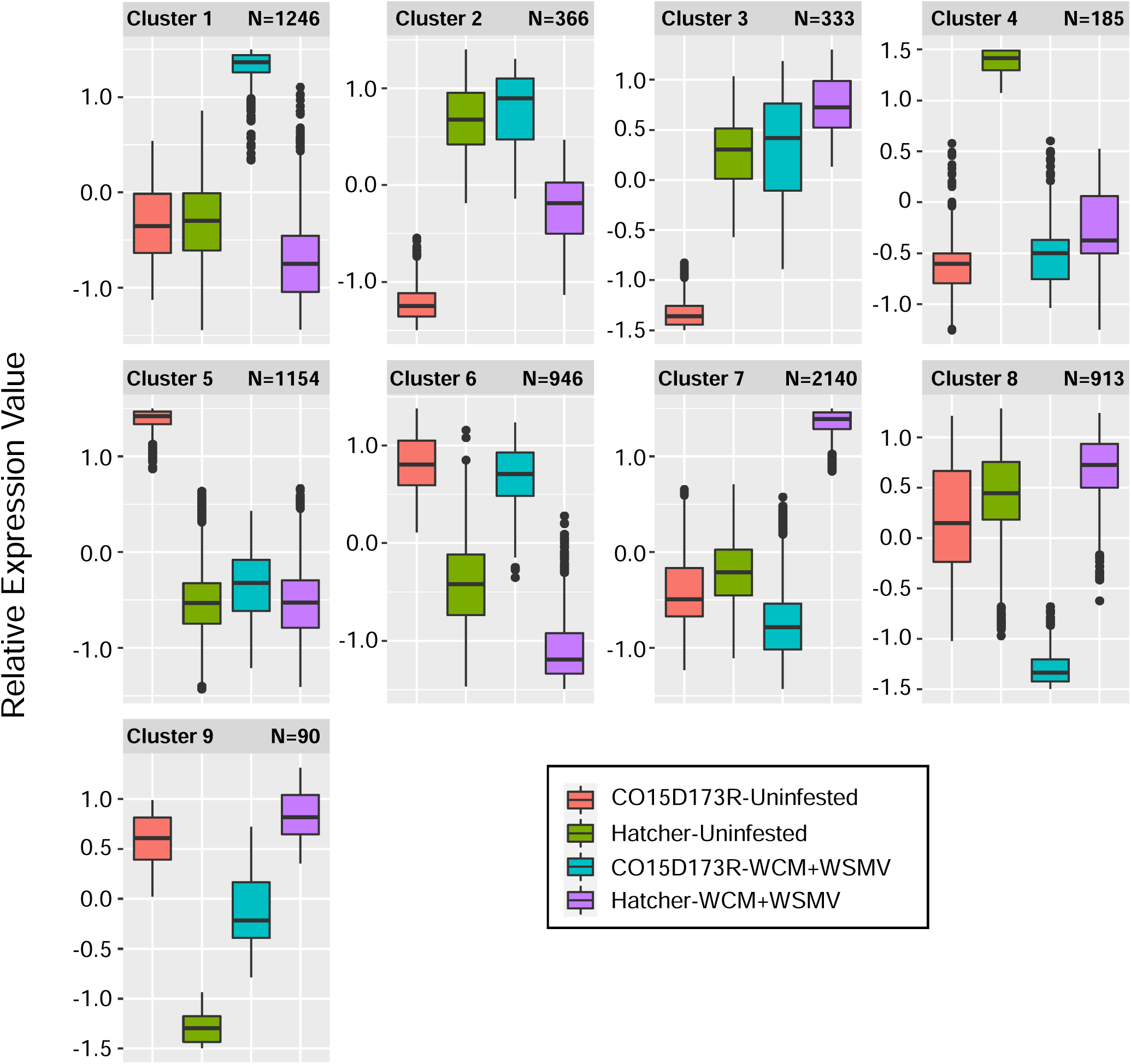
Expression profiles of the DEGs before and after WCM and WSMV infestation that are organized in 9 clusters. Boxplots represent the related gene expression values (red=CO15D173R control, green=Hatcher control, turquoise=CO15D173R infested and purple=Hatcher infested).

The clustering analysis also identified clusters in which potential defense mechanisms are present. For instance, cluster 2 consists of genes with high expression in the susceptible-infested and tolerant uninfested conditions and the functions of these genes were related to structural constituent of ribosome or structural molecule activity. Genes with a high expression level in all the conditions, except in wheat susceptible uninfested were part of the cluster 3 with functions related to protein transmembrane activity, phosphatidylethanolamine binding, oxidoreductase activity or UDP-glucose 4-epimerase activity. Genes with a high expression in the tolerant genotype and low expression in the 3 remaining conditions were found in the cluster 7 with functions related to ammonia-lyase activity, calcium ion binding, oxidoreductase activity, hormone binding or positive regulation of defense response to insect (Biological Process) or immune system (Biological Process). The cluster 8 was composed of genes with a high expression level in all the conditions except susceptible infested. The functions were related to vitamin binding, transaminase activity, oxidoreductase activity, hydrolase activity, ATPase-couples transmembrane transporter activity (BP: response to biotic stimulus, maltose biosynthetic process, small molecule catabolic process or lipid oxidation).

### Transcription factors

In total, 162 transcription factors (TFs) were differentially expressed in our dataset and were organized into 36 families with 1 to 28 genes per family (Supplemental Table 2). In total, 70 and 50 TFs were downregulated and upregulated in the wheat-tolerant genotype, respectively. In contrast, 18 and 11 were down and upregulated, respectively, in the susceptible genotype. Fourteen TFs families were composed of at least four genes (e.g., AP2 and C2H2) with a maximum of 28 genes for the NAC family. Among the 28 genes related to the NAC family, 16 were specifically upregulated in the wheat-tolerant genotype, and eight were downregulated in the wheat-tolerant genotype (Figure 8). Comparing the TF families up or downregulated in the tolerant genotype, five TF families, G2-like, bZIP, TALE, AP2, and Dof, were found to be downregulated only. Among the TFs upregulated in the susceptible or tolerant wheat genotype, MYB-related and ERF families were upregulated only in the tolerant genotype and not in the susceptible genotype. The expression level of the TFs part of the NAC families has 14 genes with an expression level higher in the tolerant genotype infested compared to the other three conditions (Figure 8).

**Figure 8:**
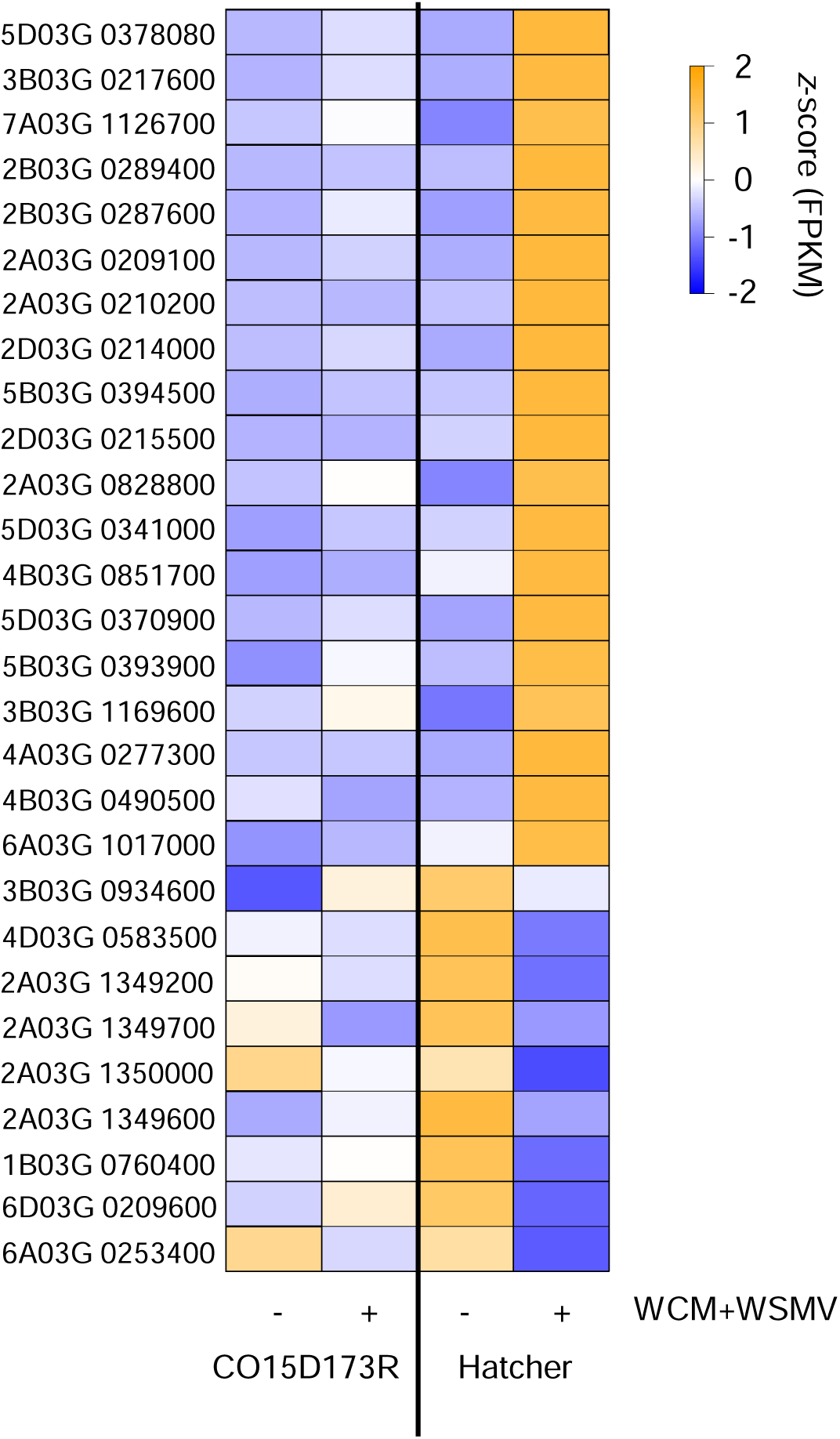
Heatmap of the expression of WCM and WSMV infestation induced change of genes encoding NAC TF family. Each column corresponds to a condition. Each cell contains the corresponding normalized FPKM value. Expression level is giving with the color scale: orange=high expression level, blue: low expression level.

## Discussion

WCM and mite-vectored viruses continue to cause significant yield losses to wheat production in the US Great Plains (Appel et al. 2015; Hollandbeck et al. 2020; Albrecht et al. 2022). In this study, we investigated the molecular basis of tolerance to this dual threat in the wheat genotype, Hatcher. Our transcriptomic analysis revealed a distinct pattern: genes involved in defense hormone pathways (JA, SA, and ABA) were upregulated, while those related to cell division and growth were downregulated, aligning with the plant growth-defense trade-off theory (He, Webster, and He 2022). Notably, we observed the upregulation of TFs associated with lignin biosynthesis, suggesting a potential role in conferring the tolerance phenotype to Hatcher. This research provides novel insights into the potential mechanism of combined WCM and WSMV tolerance in wheat, advancing our understanding beyond previous studies that focused solely on WCM or investigated transcriptomic responses at short (1 day) or prolonged (10 days) feeding durations (Kiani et al. 2021; Pingault et al. 2022).

Tolerance to WCM and WSMV has not been previously reported for any genotype of wheat plants. The consistent prevalence of WSMV and presumably WCM across the experimental site, as evidenced by both ELISA and qRT-PCR data, highlights the persistent disease pressure faced by all wheat varities throughout the study period. This consistent disease pressure underscores the validity of our findings, as it provides a robust background against which to assess varietal responses. Interestingly, despite this high and consistent disease pressure, significant yield differences were observed between varities, particularly in 2015 and 2016. Hatcher, despite showing high WSMV incidence, consistently outperformed other varities in terms of yield, even surpassing Snowmass (carrying the *Wsm2* resistance gene; ( Haley et al. 2011)) and Mace (carrying the *Wsm1* resistance gene; (Graybosch et al. 2009)). This unexpected finding suggests that Hatcher possesses unique tolerance mechanisms to combined infestation by WCM and WSMV, independent of known resistance genes. Notably, Hatcher has also demonstrated partial tolerance to another significant wheat pest, the wheat stem sawfly (WSS; *Cephus cinctus*).

Previous research (Lavergne et al. 2020) revealed distinct proteomic and metabolomic changes in Hatcher in response to WSS infestation, implicating pathways related to enzymatic detoxification, proteinase inhibition, and antiherbivory compound production. Collectively, these studies suggest a potential broader tolerance mechanism in Hatcher, extending beyond WSMV and encompassing responses to multiple biotic stressors.

### Phytohormal Regulation: A Balancing Act

Phytohormones orchestrate complex responses to biotic stress, often involving trade-offs between defense mechanisms and growth processes. In our study, this intricate balance was evident in the differential regulation of various hormone pathways. JA plays a complex role in plant defense against phloem-feeding insects, with responses varying depending on feeding duration and JA levels. In sorghum, short-term aphid feeding triggers a transient increase in JA levels that deters aphids, while prolonged feeding increases susceptibility due to elevated JA levels (Grover et al. 2022). Similarly, both low and high levels of JA have been shown to positively influence wheat’s response to aphid feeding (Aslam et al. 2022). Our findings in wheat demonstrate a similar complexity in the JA response to WCM and WSMV infestation. While genes involved in JA biosynthesis were upregulated in the tolerant genotype, JA-responsive defense genes like *JAR1* were downregulated or not differentially expressed. This suggests that increased JA levels alone may not be sufficient to activate downstream defense mechanisms against WCM and WSMV in wheat.

ABA, often associated with abiotic stress responses, has been implicated in biotic stress tolerance as well, with elevated levels contributing to soybean tolerance to soybean aphids (Chapman et al. 2018). ABA interacts with JA to mutually enhance their biosynthesis through the ABA-induced expression of *Myc2* (Luo et al. 2023). This interplay triggers a signaling cascade where ABA binds to the PYL/PYL/RCAR complex, repressing PP2Cs, allowing the expression of *SnRK2*, and activating ABF transcription factors, ultimately activating ABA-responsive genes. Our transcriptomic analysis revealed a similar pattern in wheat responding to WCM and WSMV infestation. Three *PP2C* genes were downregulated in the tolerant genotype while remaining unchanged in the susceptible genotype. Additionally, two *ABF* genes were upregulated in the tolerant genotype and downregulated in the susceptible one, despite both showing upregulation of *SnRK2* genes. These findings strongly suggest a pivotal role for ABA in mediating wheat tolerance to WCM and WSMV infestation.

Interestingly, genes responsive to auxin and SA were primarily upregulated in the tolerant genotype. This observation implies a potential compensatory mechanism where other hormonal pathways, such as auxin and SA signaling, may be more prominent in conferring WCM and WSMV tolerance in wheat. Further investigation into the interactions between JA, auxin, and SA pathways could elucidate the precise mechanisms underlying this complex defense response.

Wheat development progresses through vegetative and reproductive phases, with tiller production during the vegetative phase being a key determinant of eventual grain yield (Ying Wang, Miao, and Yan 2016). Our study observed continued head production in uninfested plants of both genotypes beyond 55 days post-inoculation (dpi). However, WCM and WSMV infestation did not significantly alter tiller numbers between 55 and 69 dpi in the tolerant Hatcher or susceptible CO15D173R genotypes. This observation may be linked to the differential regulation of phytohormones involved in growth and development. Gibberellic acid (GA), essential for stem elongation and germination (Gupta and Chakrabarty 2013), and brassinosteroids (BR), known to regulate tiller number in wheat (Shang et al. 2021), showed contrasting expression patterns. Genes involved in GA and BR biosynthesis were upregulated in the tolerant Hatcher genotype and downregulated in the susceptible genotype. This suggests a potential mechanism by which Hatcher may maintain growth-related processes even under WCM-induced stress, highlighting the complex interplay between growth regulation and stress tolerance in wheat.

### Lignin Biosynthesis: A Potential Defense Barrier

Our study revealed a significant upregulation of genes involved in lignin biosynthesis in the WCM and WSMV tolerant Hatcher, while no differential expression was observed in the susceptible genotype. This suggests a potential role for the lignin pathway in conferring tolerance against WCM and WSMV. The importance of lignin in plant defense is supported by previous findings in Arabidopsis and *Chrysanthemum morifolium*, where MYB transcription factor overexpression led to increased expression of lignin biosynthesis genes (Yinjie Wang et al. 2017). In Hatcher, we identified three MYB genes co-expressed and upregulated with lignin biosynthesis genes, further strengthening this link. Increased lignin content influences leaf morphology, potentially impacting the integrity and size of bulliform cells responsible for leaf curling (Sun et al. 2020). By reducing leaf curling, the plant may offer less refuge to WCM, thereby limiting population growth and contributing to overall tolerance.

### WSMV Resistance: A Complex Landscape

Three loci associated with WCM resistance (*Cmc2*, *Cmc3*, and *Cmc4*) have been previously identified (Malik et al. 2003; Zhao et al. 2021; Whelan and Hart 1988). However, only *Cmc4* (*TraesCS6D01G005300*) has been characterized at the genomic level (Zhao et al. 2021), and our study found no differential expression of this gene in response to WCM and WSMV infestation in either wheat genotypes. Plant viruses, including WSMV, employ various strategies to evade or suppress host defenses, such as gene silencing. In wheat, four genes (*Wsm1*, *Wsm2*, *Wsm3*, and *c2652*) have been implicated in WSMV resistance (S. d. Haley et al. 2002; Sharp et al. 2002; Divis et al. 2006; Haber, Seifers, and Thomas 2006). Notably, our analysis revealed the upregulation of two chaperone protein DnaK genes (*Traes5A03G0661300* and *TraesCS5B03G0699800*), located within the *Wsm2* locus, in the tolerant Hatcher (Xie et al. 2022). This observation reinforces the potential importance of these genes in WSMV tolerance. Furthermore, our dataset showed differential expression of 70 genes related to disease-resistance proteins. Among these, 13 genes were upregulated in Hatcher and linked to the SA signaling pathway, known to play a role in plant defense. Interestingly, nine of these genes belong to the *TaPR1* locus, previously identified as essential for stripe rust resistance in wheat and associated with lignin accumulation after fungal infection (Liu et al. 2023). This finding suggests a potential convergence of defense mechanisms against diverse pathogens in wheat, linking WSMV tolerance to specific resistance genes and broader defense responses.

### Conclusions

In conclusion, our study elucidates the mechanisms underlying the tolerance of the wheat genotype Hatcher to the combined threat of WCM and WSMV. A delicate balance between defense and growth responses is evident, with the upregulation of genes involved in JA, SA, and ABA pathways highlighting a robust defense response in the tolerant genotype. Additionally, the activation of lignin biosynthesis genes suggests a potential physical barrier against these pests. The differential regulation of genes associated with cell division and growth hormones such as GA and BR underscores their role in maintaining plant growth despite pest pressure. Overall, our findings provide valuable insights into the molecular basis of wheat tolerance to WCM-WSMV, paving the way for the development of resistant varieties and improved pest management strategies in wheat production.

## Supplementary material Data

### Availability Statement

RNA-seq data are available at https://dataview.ncbi.nlm.nih.gov/object/PRJNA998910?reviewer=kghls7f3v4009bq06e3clmjvlf. BioProject PRJNA998910.

## Acknowledgements

We would like to thank Joe Schneekloth at CSU Extension for assistance planting and harvesting field trials at Akron Co, and Dr. Ned Tisserat for completing the field trials in 2015. Lise Pingault was partially supported by the UNL Research Council: Faculty Seed Grants awarded to Lise Pingault and by US National Science Foundation CAREER Grant IOS-1845588 awarded to Joe Louis. The research was supported by funding from the Colorado Wheat Research Foundation to Punya Nachappa and Vamsi Nalam and funding from by the United States Department of Agriculture – Agricultural Research Service to Kirk Broders.

## Author Contributions

VJN, PN, JL and KB conceieved and planned the experiments. TA, JR, LP and KB carried out the experiments. LP and JR performed transcriptomic analysis. LP, KB, JL, PN and VN wrote the manuscript with input from all authors. All authors read and approved the final manuscript.

## Conflict of Interest

The authors have declared that no competing interests exist. Mention of trade names or commercial products in this publication is solely for the purpose of providing specific information and does not imply recommendation or endorsement by the U.S. Department of Agriculture. The U.S. Department of Agriculture prohibits discrimination in all its programs and activities on the basis of race, color, national origin, age, disability, and where applicable, sex, marital status, familial status, parental status, religion, sexual orientation, genetic information, political beliefs, reprisal, or because all or part of an individual’s income is derived from any public assistance program. (Not all prohibited bases apply to all programs.) Persons with disabilities who require alternative means for communication of program information (Braille, large print, audiotape, etc.) should contact USDA’s TARGET Center at (202) 720-2600 (voice and TDD). To file a complaint of discrimination, write to USDA, Director, Office of Civil Rights, 1400 Independence Avenue, S.W., Washington, D.C. 20250-9410, or call (800) 795-3272 (voice) or (202) 720-6382 (TDD). USDA is an equal opportunity provider and employer.

---

**Supplemental Figure 1:**
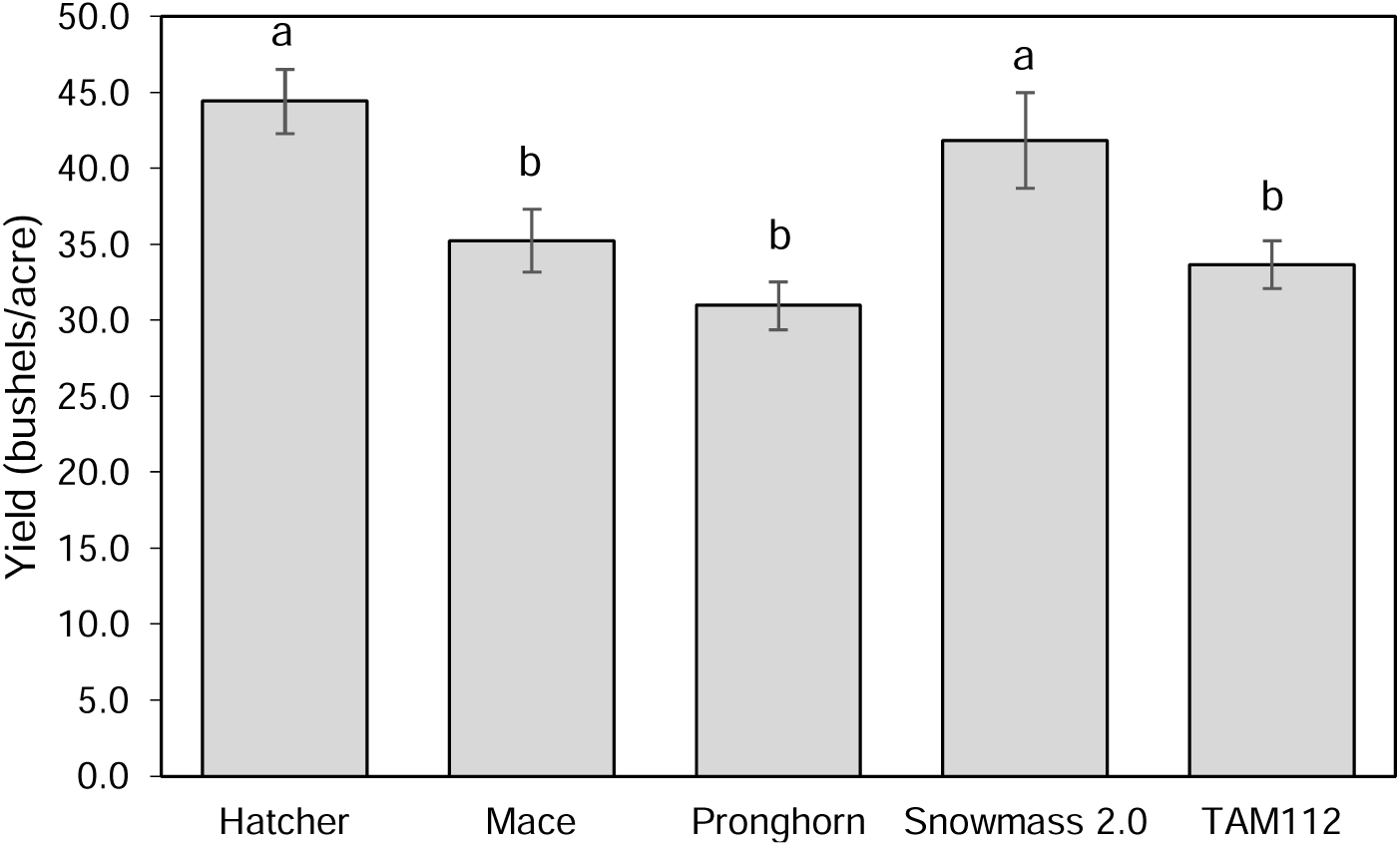
Yield averaged across the three years of the experiment. Wheat genotypes with the same letter are not significantly different (Tukey’s HSD; P<0.05).

**Supplemental Figure 2:**
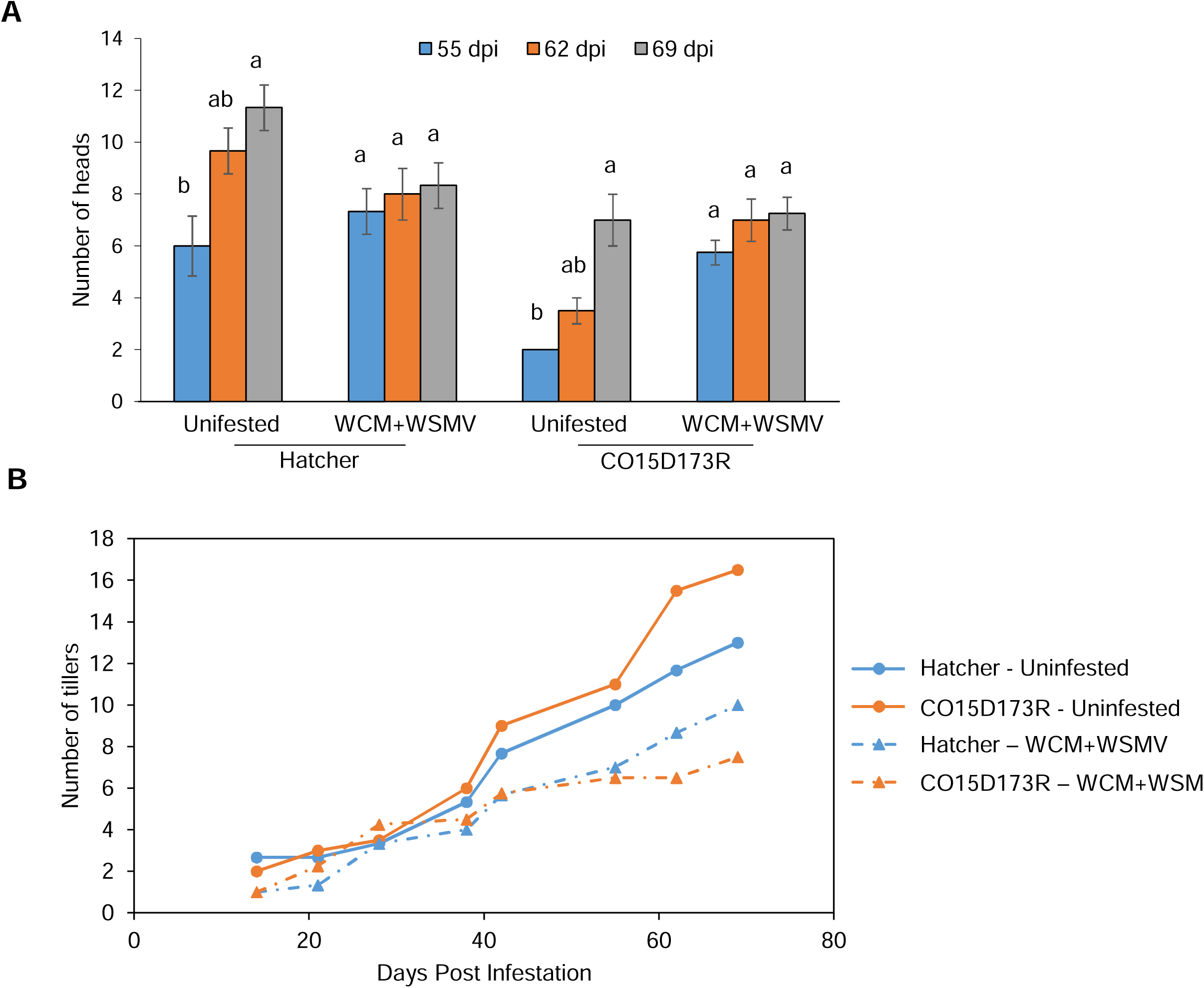
Phenotyping measurements for (A) number of heads and (B) number of tillers for CO15D173R (WCM-susceptible) and Hatcher (WCM-tolerant) during a time course. N = 3-4.

**Supplemental Figure 3:**
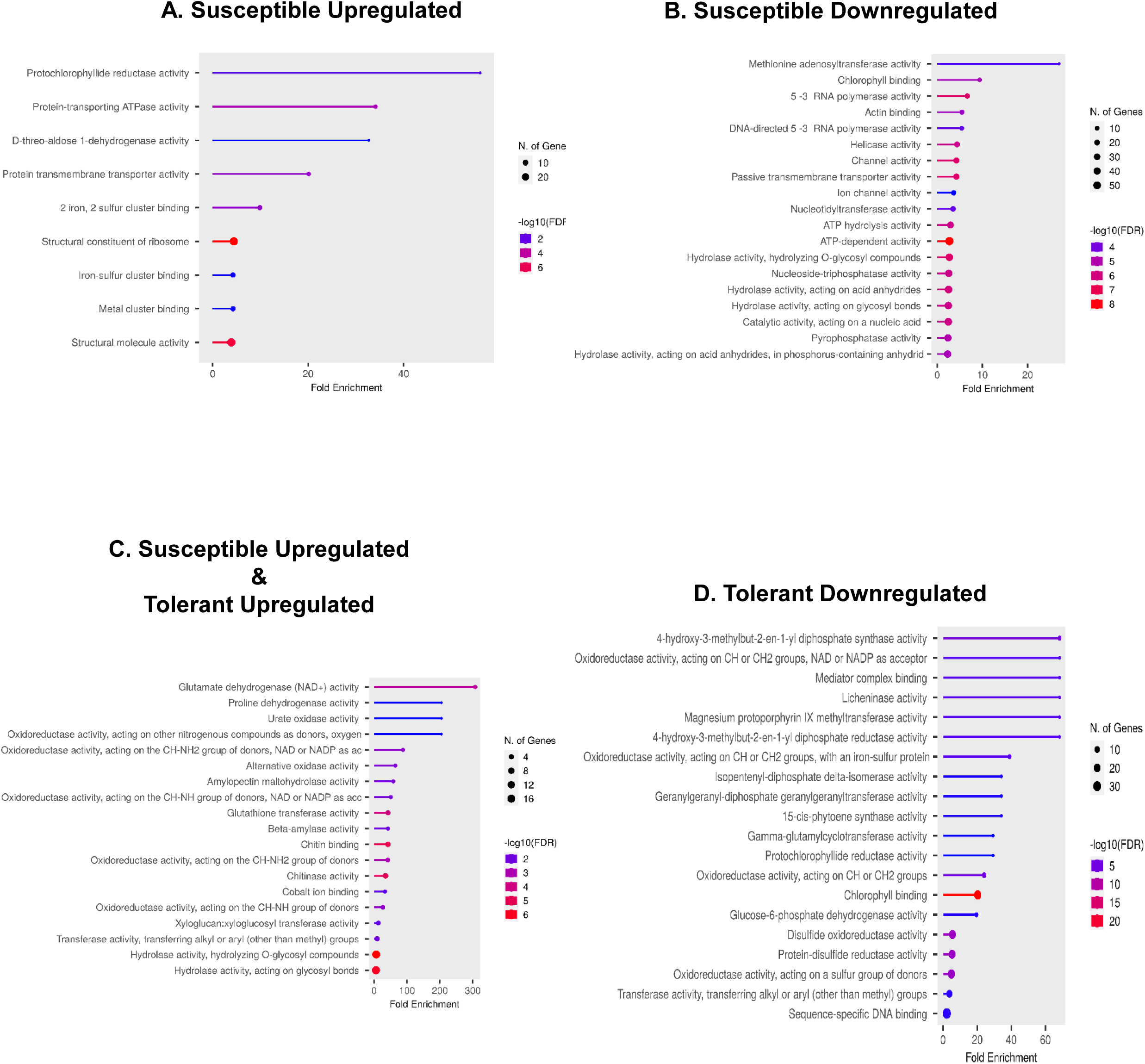
Representation of the functional enrichment for the genes (A) upregulated in the susceptible genotype, (B) downregulated in the susceptible genotype, (C) upregulated in both susceptible and tolerant genotypes and (D) downregulated in the tolerant genotype.

**Supplemental Table 1:** Mapping summary table containing number of reads sequenced and mapped on the wheat reference genome.

**Supplemental Table 2:** Gene expression contrast analysis.Highlight differentially expressed genes (DEGs), log2 fold-change values, *p*-values and FPKM values, as well as gene functions.

## Notes

### Competing Interest Statement

The authors have declared no competing interest.

